# Validation analysis of EMDB entries

**DOI:** 10.1101/2021.11.02.466942

**Authors:** Zhe Wang, Ardan Patwardhan, Gerard J. Kleywegt

## Abstract

The Electron Microscopy Data Bank (EMDB) is the central archive of the electron cryo-microscopy (cryo-EM) community for storing and disseminating volume maps and tomograms. With input from the community, EMDB has developed new resources for validation of cryo-EM structures, focussing on the quality of the volume data alone and that of the fit of any models, themselves archived in the Protein Data Bank (PDB), to the volume data. Based on recommendations from community experts, the validation resources are developed in a three-tiered system. Tier 1 covers an extensive and evolving set of validation metrics, including tried and tested as well as more experimental ones, which are calculated for all EMDB entries and presented in the Validation Analysis (VA) web resource. This system is particularly useful for cryo-EM experts, both to validate individual structures and to assess the utility of new validation metrics. Tier 2 comprises a subset of the validation metrics covered by the VA resource that have been subjected to extensive testing and are considered to be useful for specialists as well as non-specialists. These metrics are presented on the entry-specific web pages for the entire archive on the EMDB website. As more experience is gained with the metrics included in the VA resource, it is expected that consensus will emerge in the community regarding a subset that is suitable for inclusion in the tier 2 system. Tier 3, finally, consists of the validation reports and servers that are produced by the Worldwide Protein Data Bank (wwPDB) Consortium. Successful metrics from tier 2 will be proposed for inclusion in the wwPDB validation pipeline and reports. We describe the details of the new resource, with an emphasis on the tier 1 system. The output of all three tiers is publicly available, either through the EMDB website (tiers 1 and 2) or through the wwPDB ftp sites (tier 3), although the content of all three will evolve over time (fastest for tier 1 and slowest for tier 3). It is our hope that these validation resources will help the cryo-EM community to get a better understanding of the quality, and the best ways to assess the quality of cryo-EM structures in EMDB and PDB.

## 1. Introduction

Structural biology has been revolutionised by cryogenic electron microscopy techniques (cryo-EM), that produce Coulomb potential maps of biomolecules, complexes and assemblies that have often proved difficult to resolve by X-ray crystallography (Kühlbrandt, 2014). Improvements in microscopy, detection and computational methods have in favourable cases even enabled atomic-resolution (∼1.2Å) structure determination (Yip *et al*., 2020; Nakane *et al*., 2020). These developments have led to a rapid increase in the number of cryo-EM structures being determined and deposited in the Electron Microscopy Data Bank (EMDB; Tagari *et al*., 2002; Lawson *et al*., 2016) as shown in **Figure 1**, and in many cases allow for atomic models to be built and deposited in the Protein Data Bank (PDB; wwPDB Consortium, 2019). While single-particle averaging (SPA) methods provide the bulk of these structures, there is an increasing interest in structure determination *in situ*. Here, multiple copies of an object of interest (e.g., ribosomes, virus particles or nuclear pore complexes) are identified in and extracted from cryogenic electron tomograms (cryo-ET), and then improved in an iterative process of superposition and averaging (Briggs, 2013; Schur *et al*., 2015).

**Figure 1.**
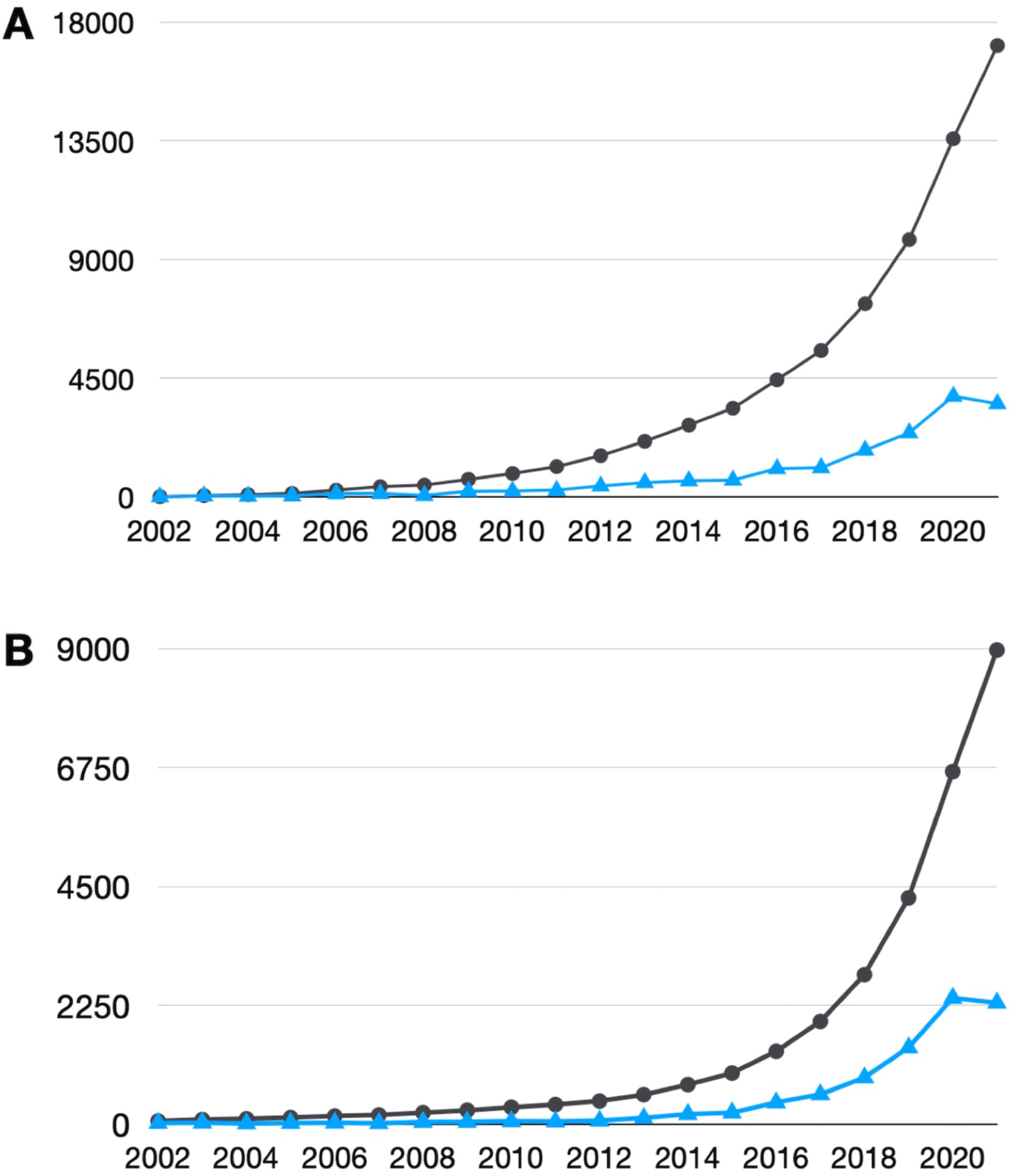
Trends in archiving of cryo-EM structures. (A) The number of released EMDB entries per year (blue) and the cumulative number of entries (black), as of October 2021 (data from: https://www.ebi.ac.uk/emdb/statistics). (B) The annual (blue) and cumulative (black) number of EM-based structures in the PDB as a function of year, as of October 2021 (data from: https://www.rcsb.org/stats/growth/growth-em).

As always with structure determination based on experimental data, there is a need to validate the data, the model constructed based on the data, and the fit of model and data (Kleywegt, 2000; Gore et al., 2017). Fortunately, the cryo-EM field can build on the wide experience with validation methods gained by protein crystallographers over the past three decades, which has resulted in community-wide agreement regarding sensible validation checks to apply to new X-ray structures upon deposition to the PDB (Read *et al*., 2011). Much of the software and methods developed, in particular for model validation (e.g., O (Jones *et al*., 1991), WHAT_CHECK (Hooft *et al*., 1996), MolProbity (Davis *et al*., 2007; Williams *et al*., 2018)), can also be used for models derived by other methods such as cryo-EM (Henderson *et al*., 2012) and NMR spectroscopy (Montelione *et al*., 2013). Such methods assess adherence of models to known geometric, physical, stereo-chemical and conformational criteria (e.g., bond lengths and angles, non-bonded distances, chirality of amino acids and nucleic acids, and preferred combinations of main-chain and side-chain torsion angles). However, because of the different nature of the underlying experimental data, methods for data validation and model/data-fit assessment in cryo-EM have to be developed from scratch, although some methods used in the X-ray field can be adapted (e.g., various residue-based model/data-fit criteria; Lawson *et al*., 2021). In cryo-EM, the raw experimental data is only available for a minority of structures (in the EMPIAR archive; Iudin *et al*., 2016) and there are no established methods to assess a model’s fit to the underpinning raw data. Hence, in practice the quality is assessed instead of the map derived from the raw data as well as of the fit between the model and that map.

EMDB and the Worldwide Protein Data Bank Consortium (wwPDB; Berman *et al*., 2003) established a Validation Task Force for cryo-EM (EM-VTF) in 2010 which published its recommendations two years later (Henderson *et al*., 2012). Inspired by this, EMDB developed so-called Visual Analysis pages for every EMDB entry (Lagerstedt *et al*., 2013). These pages contained such elements as orthogonal projections of maps, orthogonal surface views of maps and models, atom-inclusion plot, FSC curve, etc. However, both the first EM-VTF meeting and the Visual Analysis functionality predated the “resolution revolution” (Kühlbrandt, 2014) and with the rapidly increasing number of moderate to high-resolution cryo-EM structures being determined there was a need to reassess the state of the field and to update the recommendations regarding archiving and validating these structures. This happened at the wwPDB single-particle cryo-EM data-management workshop in early 2020 (Kleywegt *et al*., to be published), again organised jointly by EMDB and wwPDB. This meeting resulted in a large number of recommendations regarding the validation criteria that should be used by the archives, and also identified a number of areas where there is no community agreement yet regarding which method or software is best suited to validate a certain aspect of the map or map/model fit (Kleywegt *et al*., to be published).

Prior to the wwPDB single-particle cryo-EM data-management workshop, refactoring of the Visual Analysis functionality had begun to use up-to-date technologies and add some functionality. The new system is referred to as the Validation Analysis (VA) resource (https://emdb-empiar.org/va), and it continues to present its results on a single webpage for each EMDB entry. Following the recommendations of the 2020 workshop, we have developed a three-tiered strategy to implement, test and disseminate validation results on cryo-EM structures (with or without a model): a development version (tier 1, the VA resource), a production version (tier 2, part of the EMDB entry pages) and a version incorporated into the wwPDB validation-pipeline (tier 3).

The VA resource (tier 1) is the full development version, aimed at specialist users, offering a rich and evolving selection of validation data. New validation methods are implemented here first and run on the entire archive. This enables archive-wide analyses to assess the usefulness, robustness, reliability, etc., of these methods and allows individual specialists to assess how they perform in specific cases. An example URL for a VA page (for EMDB entry EMD-11145; Toelzer *et al*., 2020) is: https://www.ebi.ac.uk/emdb/va/EMD-11145. Once community consensus has been reached regarding the utility of certain validation criteria, these can be added to the production version (tier 2). On the other hand, if there are questions regarding the soundness of any criterion, we may either continue to expose it in tier 1 to give the community more time to experiment with and assess it, or if the concerns are major, we may drop it from tier 1 altogether.

Tier 2 is a scaled-down version of the VA resource containing validation components that are well-tested and generally considered valuable, robust, informative and well-understood. These pages are accessible in a separate tab (labelled Validation) of the EMDB entry pages for each EMDB structure. The tier 2 page for the same example entry as before can be found at: https://www.ebi.ac.uk/emdb/EMD-11145?tab=validation.

Finally, following agreement with wwPDB (as of 1 January 2021 EMDB is a part of the wwPDB Consortium), some of the most informative criteria will be implemented in tier 3, the validation pipeline that is part of the OneDep deposition, annotation and validation system (Young *et al*., 2017) and its validation servers. Any validation components in tier 3 are thus applied to every newly deposited cryo-EM volume in EMDB and accompanying model in the PDB, and these reports are also made available for the entire EMDB archive and all EM structures in the PDB. Many of the recommendations from the wwPDB single-particle cryo-EM data-management workshop have already been implemented in one or more of the validation tiers.

Here, we describe the current state of the VA resource (tier 1). Some of its elements are also part of the production version (tier 2) or even the OneDep validation pipeline (tier 3).

## 2. Validation Analysis resource

It is useful to understand what types of data may be archived as part of a single EMDB entry; this is summarised in **Table 1**.

**Table 1.** Description of data types that can be deposited to EMDB.

| Category (number) | Explanation or examples |
| --- | --- |
| Primary map or tomogram (or tomogram average) (1) - mandatory | EM map or tomogram that is shown and described in the associated literature publication |
| Raw map (0-1) * | Unfiltered, unsharpened and unmasked raw map on which the primary map is based |
| Half-maps (0 or 2) ** | Unfiltered, unsharpened and unmasked raw half-maps for single-particle analysis, single-particle-based helical reconstructions or sub-tomogram averaging |
| Masks (0 or more) | Primary/raw map masks, segmentation/focused refinement masks and half-map masks |
| Segmentations (0 or more) *** | Segmentations of the map or tomogram (3D regions-of-interest; note that *masks* are computational tools whereas *segmentations* embody interpretation in terms of biological entities/objects) |
| Additional cryo-EM maps (0 or more) | Examples include difference maps, maps showing alternative conformations and/or compositions, maps with different processing (e.g., filtering, sharpening and masking) |
| Auxiliary files (0 or more) | Examples include author-determined FSC curves (half-maps, map-model, ...), structure factors and layer lines |

\* Raw maps can currently be deposited as unspecified “additional maps”. In the future it will become possible to deposit them as a separate type of map.
\*\* In the future, deposition of half-maps will become mandatory if they were used to estimate resolution.
\*\*\* This will become possible in the future.

EMDB accommodates several cryo-EM modalities including SPA, tomography, sub-tomogram averaging, helical reconstruction and electron diffraction. Any accompanying models must be deposited to the PDB while raw data (micrographs, particle stacks, tilt series, etc.) can optionally be deposited to EMPIAR (Iudin *et al*., 2016). The exact information shown in the VA resource for a specific entry depends on the modality and the presence or not of one or more models, masks and segmentations. Note that pure model validation is not included; for this users are referred to the wwPDB validation reports (Gore *et al*., 2017).

The VA page for an entry in the most general case (SPA, with a model, half-maps, masks, etc.) will include the sections shown as examples in **Figure 2**. The full contents are discussed in more detail below. **Table 2** summarises which validation components are currently implemented in each of the three tiers.

**Table 2.**
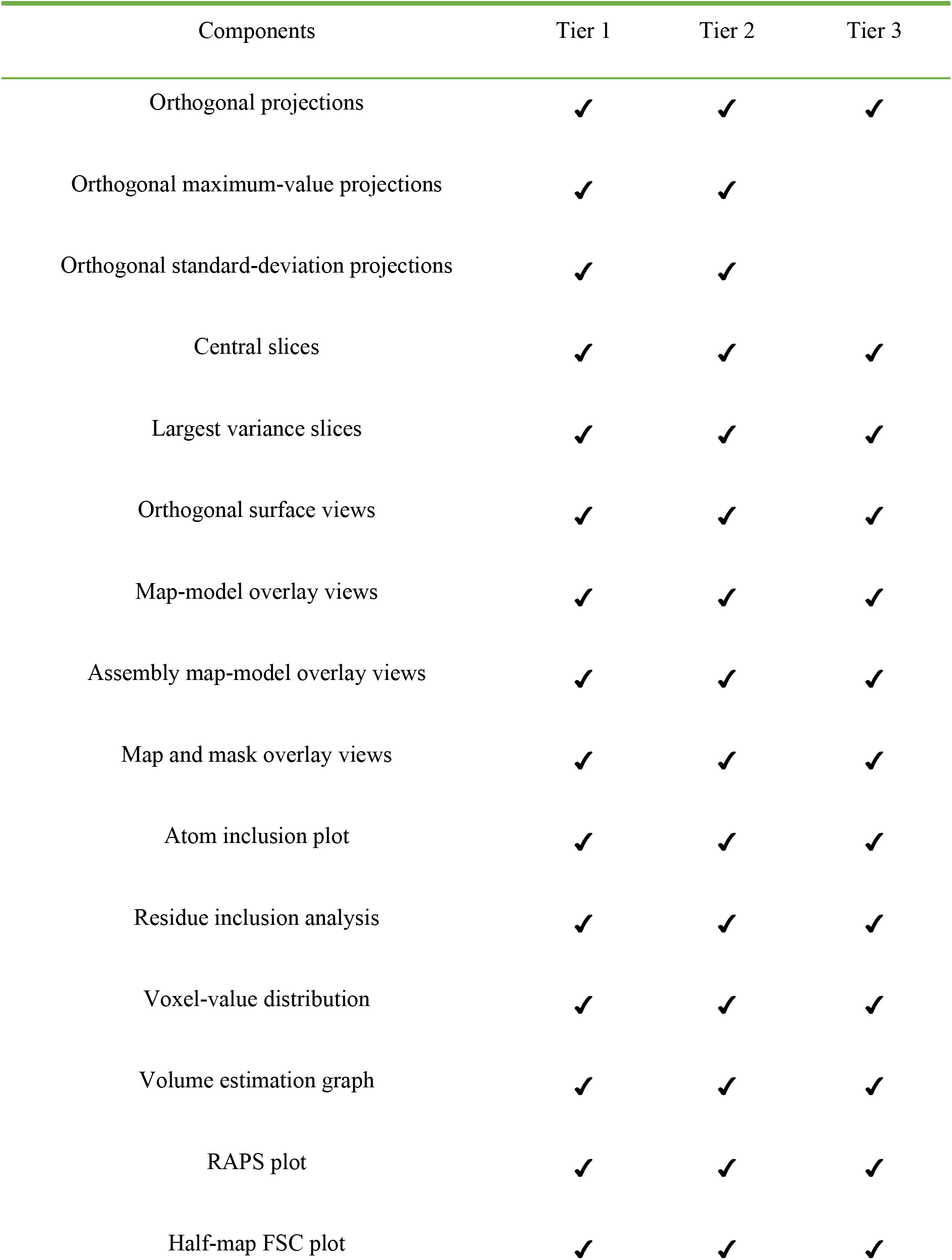
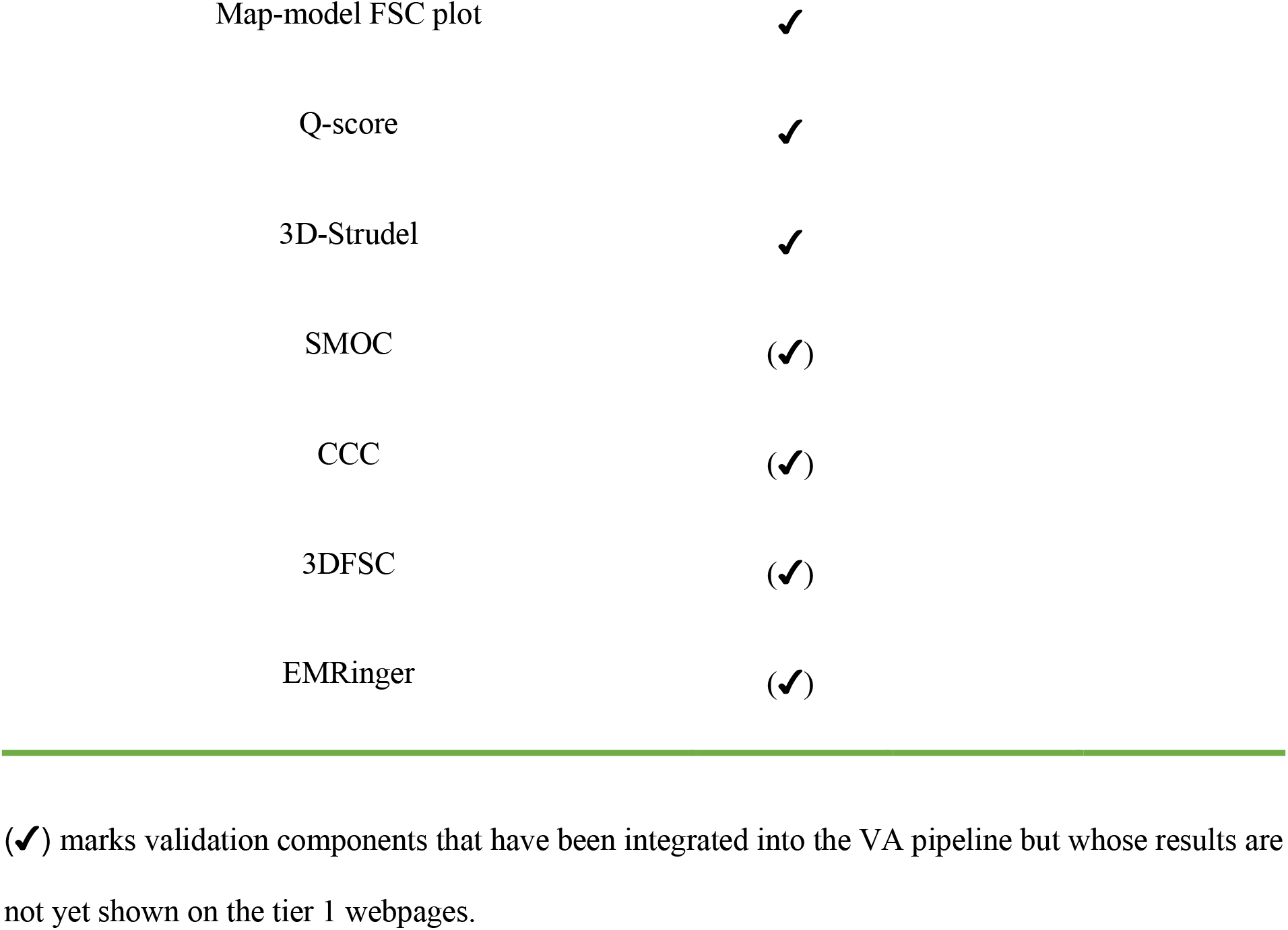
Validation components available as part of the three tiers of the EMDB validation resource (see text for details). Note that the tier 3 functionality is a subset of the tier 2 functionality, which in turn is a subset of the tier 1 components.

| Components | Tier 1 | Tier 2 | Tier 3 |
| --- | --- | --- | --- |
| Orthogonal projections | ✓ | ✓ | ✓ |
| Orthogonal maximum-value projections | ✓ | ✓ |  |
| Orthogonal standard-deviation projections | ✓ | ✓ |  |
| Central slices | ✓ | ✓ | ✓ |
| Largest variance slices | ✓ | ✓ | ✓ |
| Orthogonal surface views | ✓ | ✓ | ✓ |
| Map-model overlay views | ✓ | ✓ | ✓ |
| Assembly map-model overlay views | ✓ | ✓ | ✓ |
| Map and mask overlay views | ✓ | ✓ | ✓ |
| Atom inclusion plot | ✓ | ✓ | ✓ |
| Residue inclusion analysis | ✓ | ✓ | ✓ |
| Voxel-value distribution | ✓ | ✓ | ✓ |
| Volume estimation graph | ✓ | ✓ | ✓ |
| RAPS plot | ✓ | ✓ | ✓ |
| Half-map FSC plot | ✓ | ✓ | ✓ |

|  |  |
| --- | --- |
| Map-model FSC plot | ✓ |
| Q-score | ✓ |
| 3D-Strudel | ✓ |
| SMOC | (✓) |
| CCC | (✓) |
| 3DFSC | (✓) |
| EMRinger | (✓) |

(✓) marks validation components that have been integrated into the VA pipeline but whose results are not yet shown on the tier 1 webpages.

**Figure 2.**
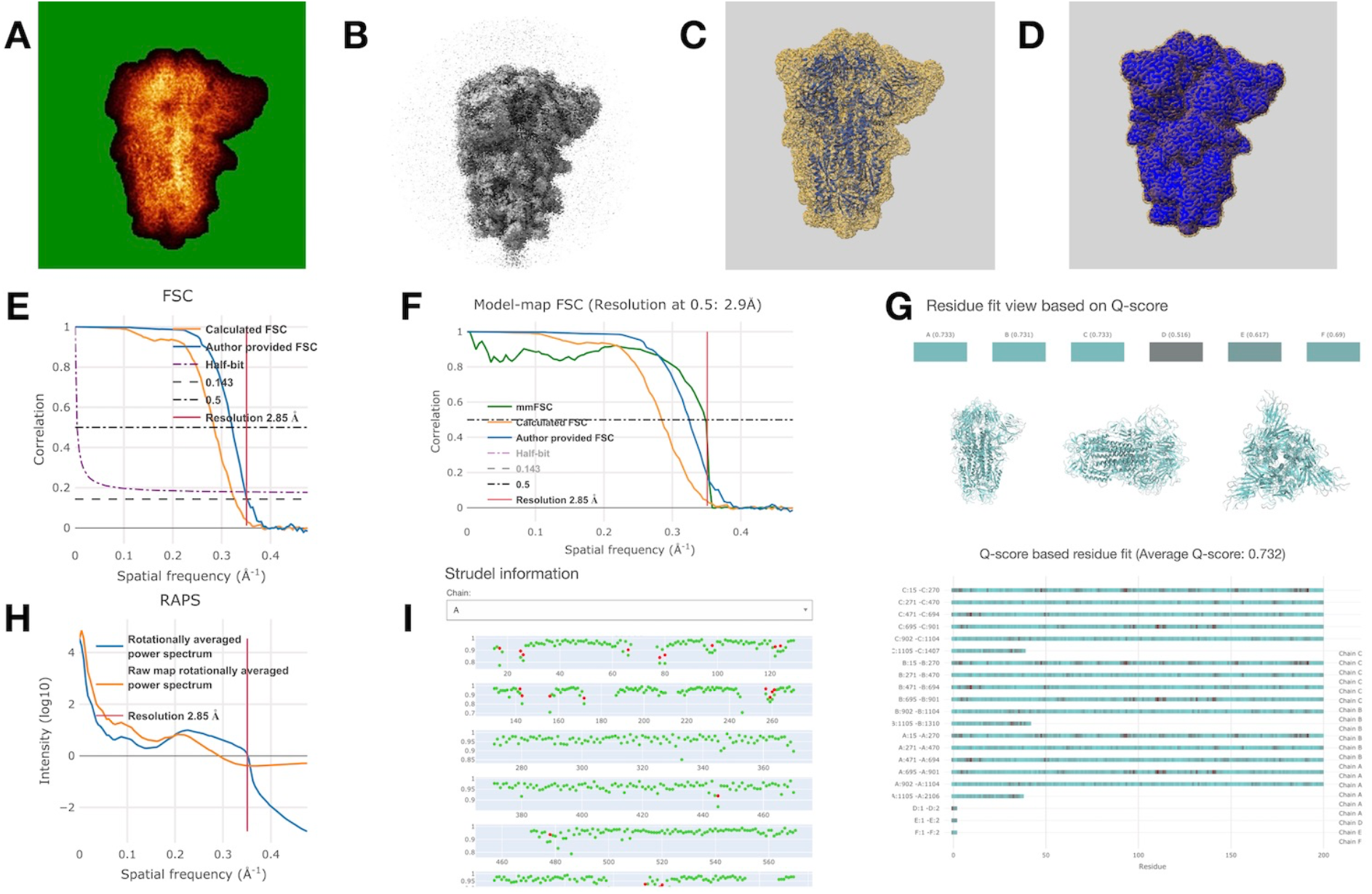
Examples of validation components in VA pages for EMDB entries (see text for details). The examples are for EMDB entry EMD-11145 (https://www.ebi.ac.uk/emdb/EMD-11145) which is a SARS-CoV-2 spike protein (Toelzer *et al*., 2020). (A) Orthogonal standard-deviation projection (shown with false colour) along the X-axis. The lowest values are mapped to solid green, which may help identify if and how masking was applied. (B) Orthogonal surface view of the raw map (calculated by averaging the two half-maps) along the X-axis. (C) Overlay of the primary map (yellow, semi-transparent) and fitted model (blue) viewed along the X-axis. (D) Overlay of the primary map (yellow, semi-transparent) with a mask (blue) viewed along the X-axis. (E) FSC plot combining the author-provided curve (cyan) and the one calculated from the two half-maps (orange) as well as various criteria used by the community to estimate resolution. The red vertical line indicates the resolution reported by the author. (F) Map-model FSC plot. FSC values are calculated between the primary map and a map calculated from the model using REFMAC (Murshudov *et al*., 1997). (G) Overview of map/model-fit analysis using the Q-score (Pintilie *et al*., 2020) both on a per-residue basis (3D view and lower part of the plot) and averaged per chain (coloured blocks at the top). The overall average score is shown in the legend (0.732 in this case). Colour-ramping is used, with cyan indicating higher Q-scores (better fit). (H) RAPS plot of the primary (cyan) and raw (orange) maps. The red vertical line indicates the author-provided resolution of the map. (I) Overview of map/model-fit analysis using 3D-Strudel (Istrate *et al*., to be published). Each row of the plot shows the 3D-Strudel score for up to 100 residues. Red dots indicate residues where the map motif of a different amino-acid type than the one in the model is most similar to the local map.

### (1) 3D volume analysis

a. *2D slices through and projections of the 3D volume*. This section allows inspection of internal details of a map or tomogram and may also be useful for identifying any artefacts (e.g., streaking or “imprints” of masks used during processing). All slices and projections are calculated and shown along the orthogonal X, Y and Z axes. In addition to standardised views of 300×300 pixels (1200×1200 for tomograms), versions with the native pixel dimensions are also provided (accessible by clicking on the standardised images). Images included are:
  i. central slices, showing the central plane in each direction.
  ii. largest-variance slices, showing the plane (in each direction) whose voxel values have the highest variance. The rationale for this is that these might be “interesting” planes.
  iii. orthogonal projections, obtained by averaging voxel values along one axis for all pixels in the plane orthogonal to that axis (e.g., the average value of all voxels along Z for each pixel in the X, Y plane).
  iv. standard-deviation projections, calculated in the same way, but showing the standard deviation rather than the average value along each projection axis.
  v. maximum-value projections, calculated in the same way, but showing the maximum value along each projection axis. All images are shown in grayscale rendering, but the maximum-value and standard-deviation projections are also shown using false colour using a slightly modified “glow” look-up table taken from Fiji (Schindelin *et al*., 2012; Schneider *et al*., 2012). This occasionally reveals features that are impossible or difficult to assess from a grayscale image (e.g., “ghosts” of masks used during processing). If half-maps are available, a raw map will be computed by averaging them, and several of the above images will also be provided for that map. To make these images comparable to those derived from the primary map, the raw map is scaled to have the same average value and standard deviation.
b. *3D views*. Except for tomograms, where there is no sensible contour level, a number of orthogonal surface views (along the X, Y and Z axes) are generated using ChimeraX (Goddard *et al*., 2018; Pettersen *et al*., 2021), including:
  i. surface of the primary map, using the recommended (usually, by the depositor) contour level.
  ii. surface of the raw map. As there is no recommended contour level for the computed raw map, it will be approximated as the level at which its surface encompasses the same volume as the primary map does at its recommended contour level. Although not always perfect, this provides a simple and consistent choice of contour level.
  iii. surface views of each mask superimposed on the primary map (which is shown as a semi-transparent surface). In the future, EMDB will add support for segmentations and these will then be shown in the same fashion as masks.
c. *Plots and graphs*. Some plots and graphs are calculated both for maps and tomograms whereas others can only be meaningfully computed for maps. They include:
  i. voxel-value histogram. The range of voxel values in a map or tomogram is divided into 128 bins and the number of voxels in each bin is plotted on a log scale. A high peak near zero is indicative of masking. If the raw map is available (see above), its voxel-value distribution will be shown in the same plot.
  ii. enclosed-volume estimate. The volume of space enclosed by a map (in nm^3^) is plotted as a function of contour level. The volume of the imaged object is estimated using the recommended contour level (shown as a vertical line). If an estimate of the molecular weight (MW) is available, then the predicted volume (using an average density of 1.5 g/cm^3^) is also indicated (by a horizontal line). Ideally, the curve and both lines intersect but there can be many causes for this not being the case, e.g., the MW may have been provided for a single unit which is repeated many times in the sample, or conversely for a larger unit than is in the map (e.g., an entire fibre). Other factors that can contribute include staining, heavier or lighter than average material in the sample, or an inaccurate contour level.
  iii. rotationally-averaged power spectrum (RAPS; Crowther & Amos, 1971; Rosenthal & Henderson, 2003) of the primary and raw maps (if available), i.e. the distribution of intensity (on a log-scale) versus spatial frequency. This may provide insight into the data-processing steps leading to the deposited primary map, in terms of CTF (contrast-transfer function) correction, low or high-pass filtering, masking artefacts and temperature-factor corrections.
  iv. FSC curves. The FSC curve between two maps calculated from independently processed half-data sets is often used to provide an estimate of the resolution limit to which both maps are still correlated (Saxton & Baumeister, 1982; van Heel & Stöffler-Meilicke, 1985; Harauz & van Heel, 1986). Here, the FSC curve calculated from the half-maps is plotted as well as the one provided by the authors (if available). Curves for several resolution-estimation criteria are also shown, as is the author-provided resolution estimate.
d. *symmetry analysis*. Symmetry analysis of a map can be useful to check that user-supplied point-group symmetry information is correct and that standard symmetry conventions for different point groups have been correctly followed. The VA resource uses ProSHADE (Nicholls *et al*., 2018) for this purpose. It produces a list of symmetries detected in the map, including the symmetry axes and a score (values smaller than ∼0.95 typically indicate partial or pseudo-symmetry), a list of all symmetry elements, and finally a list of alternative symmetries (often subgroups of the main symmetries). If symmetry was applied (and reported) this will be shown for comparison.

### (2) Map plus model assessment

There may be one, multiple or no PDB entries associated with an EMDB entry. If there are one or more models, the sections below are provided separately for each of them.

a. *3D views*. The model is shown as a blue ribbon superimposed on a semi-transparent rendering of the primary map, and viewed along the X, Y and Z axes. If point-symmetry information is available for a model (e.g., for viruses), a fully assembled model will also be generated and shown separately, also overlaid on the primary map.
b. *Atom inclusion*. An atom-inclusion graph shows the fraction of atoms that lie entirely inside the map as a function of contour level (separately for all atoms and just the backbone atoms). This plot may reveal if an inappropriate contour level was selected, or if the side chains tend not to be well contained in the map, e.g., in a low-resolution study.
c. *Map/model-fit analyses*. At present this includes information about residue inclusion (Lagerstedt *et al*., 2013), Q-score (Pintilie *et al*., 2020), 3D-Strudel (Istrate *et al*., to be published) and model-map FSC (van Heel *et al*., 2000), but this section will be expanded considerably with the inclusion (in 2022) of additional methods such as EMringer (Barad *et al*., 2015), SMOC (Joseph *et al*., 2016), CCC (Warshamanage *et al*., 2021) and 3DFSC (Tan *et al*., 2017).
  i. Residue-inclusion analysis (Lagerstedt *et al*., 2013). This is similar to the atom-inclusion metric discussed above but calculated on a per-residue basis (amino-acid residue, nucleotide or ligand). An interactive viewing panel shows the score for every residue in each molecule (in batches of 200 residues in case of longer chains). The colour of each residue depends on its inclusion score and varies from red (zero inclusion) to cyan (all atoms inside the map at the recommended contour level). In addition, three orthogonal views of the model, coloured the same way as in the panel, are shown. A significant weakness of atom and residue-inclusion scores is that they depend directly on the selected contour level. This can be chosen to be unrealistically low so as to obtain high inclusion scores, which is clearly undesirable.
  ii. Q-score (Pintilie *et al*., 2020). A more recent method, which has been calibrated using high-resolution structures and does not require any subjective parameter choices is the Q-score, a quantitative metric of resolvability in a cryo-EM map. It is calculated for individual atoms and then averaged, e.g., for a residue, molecule or complex. The presentation of the Q-score analysis is identical to that of the residue-inclusion analysis.
  iii. Map-model FSC (van Heel *et al*., 2000). The Fourier-shell correlation is calculated and plotted between the primary map and a simulated map based on the model. The graph shows to which resolution limit the model explains the map (using a cut-off value of 0.5 since the model is not independent of the map).
  iv. 3D-Strudel score (Istrate *et al*., to be published). 3D-Strudel is a model-based map-validation method which entails calculating the linear correlation coefficient between an amino-acid and rotamer-specific map-motif from the 3D-Strudel library and the cryo-EM map values around a residue of interest (after optimal superposition). The 3D-Strudel motif library was mined from EMDB maps by averaging large numbers of map fragments that correspond to each specific amino-acid rotamer in a number of resolution bands.

As explained in the introduction, the validation resources will evolve and will contain different components depending on the imaging modality and the tier. New components will be added to tier 1 (the EMDB VA resource) and, after extensive testing and evaluation by experts, some of them will be incorporated into tier 2 (EMDB entry pages). Finally, following broad community consensus and approval by wwPDB, some metrics will be included in tier 3, i.e., the wwPDB validation software pipeline, reports and servers. In the case of tier 1, some components may also be removed if they turn out to be less useful, informative or consistent, or if they provide information that is essentially redundant with other methods already incorporated.

Tier 1, 2 and 3 validation information is made available for every cryo-EM structure upon its release in EMDB (and PDB for models) and is therefore always complete and up-to-date. To reflect the evolving functionality of tier 1, its pipeline is occasionally re-run on the entire EMDB archive to make the information for all entries up to date.

## 3. Example applications

Validation-analysis components can play different roles in validating maps and any fitted models. Below we discuss some examples of issues that can be detected and diagnosed. Ideally, such issues should be addressed before deposition takes place. The depositors receive, and must approve, a preliminary validation report before being able to submit, and they are strongly recommended to run the wwPDB validation server before even commencing a deposition. The validation reports are also used by wwPDB biocurators to check entries and flag issues to the depositor prior to public release. Despite the best efforts occasionally an issue slips through the net into the public archive. The VA resource then plays a key role for EMDB staff and the cryo-EM community to check entries and rectify them after release. A few examples of this are discussed below.

**Figure 3** shows how the use of hard masks can result in artefacts. The thick green edges of the boxes apparent in the projection images show that a hard mask was applied in the periphery of the box.

**Figure 3.**
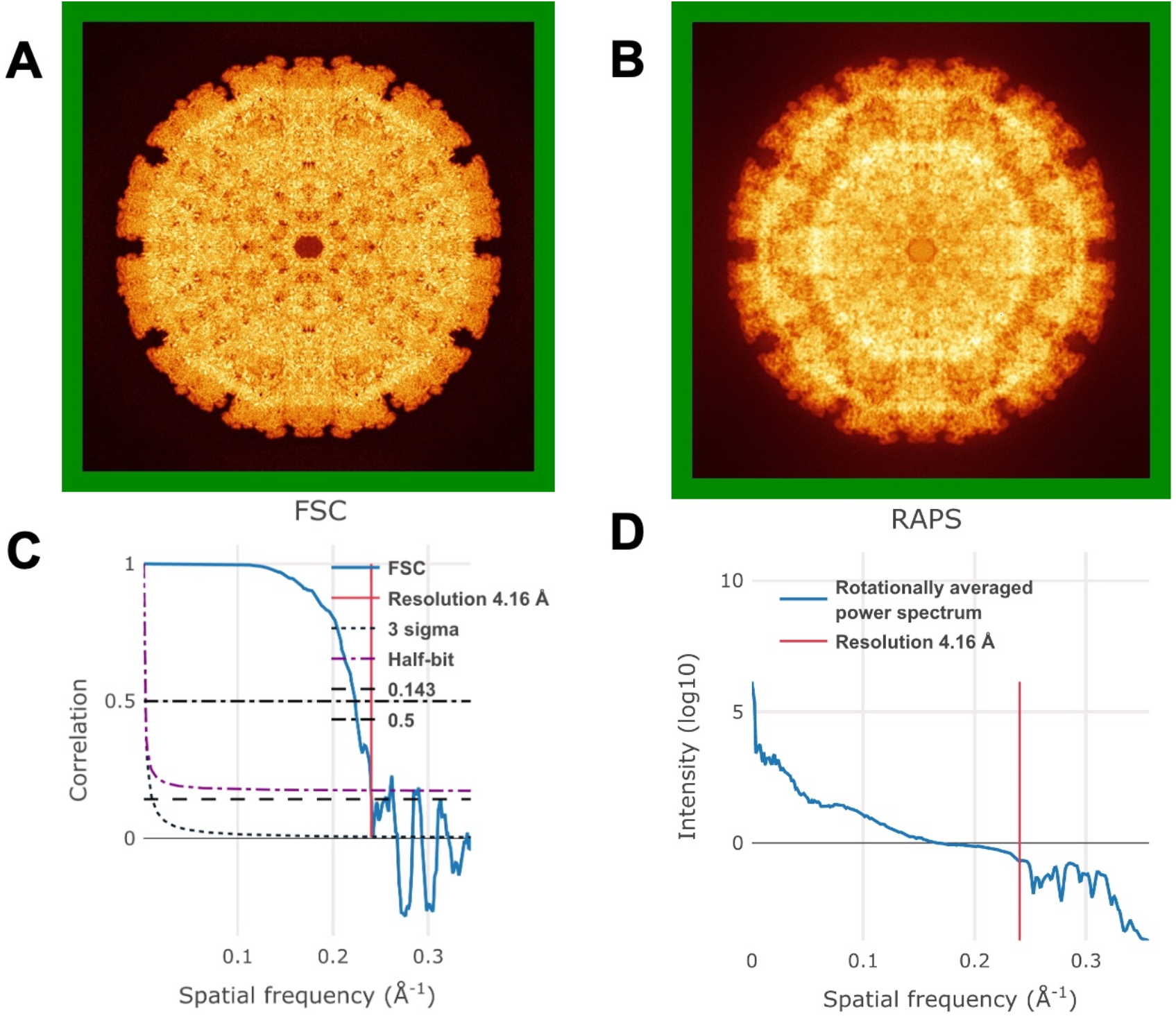
Examples of artefacts introduced due to using a hard mask (see text for details). (A) Maximum-value projection along the X-axis of a map (shown using false colour). (B) Standard-deviation projection along the X-axis of the same map (also using false colour). (C) FSC plot and (D) RAPS plot of the same map, revealing unexpected oscillations beyond the resolution cut-off.

Although the maps show good internal details, the sharp edges lead to artefacts in the FSC and RAPS plots (correlation and intensity oscillations, respectively).

If there is a fitted model, the various visualisations of map and model will provide some understanding of how well the model explains the map, allowing several fairly trivial issues to be identified and diagnosed. **Figure 4** shows an example where the map and the model are mis-aligned. The problem is easy to identify visually from the orthogonal map/model views and the model obviously shows zero atom-inclusion. Sometimes the issues are more subtle than in this case, but even then, the overlaid images and other map/model analyses often facilitate their detection and diagnosis.

**Figure 4.**
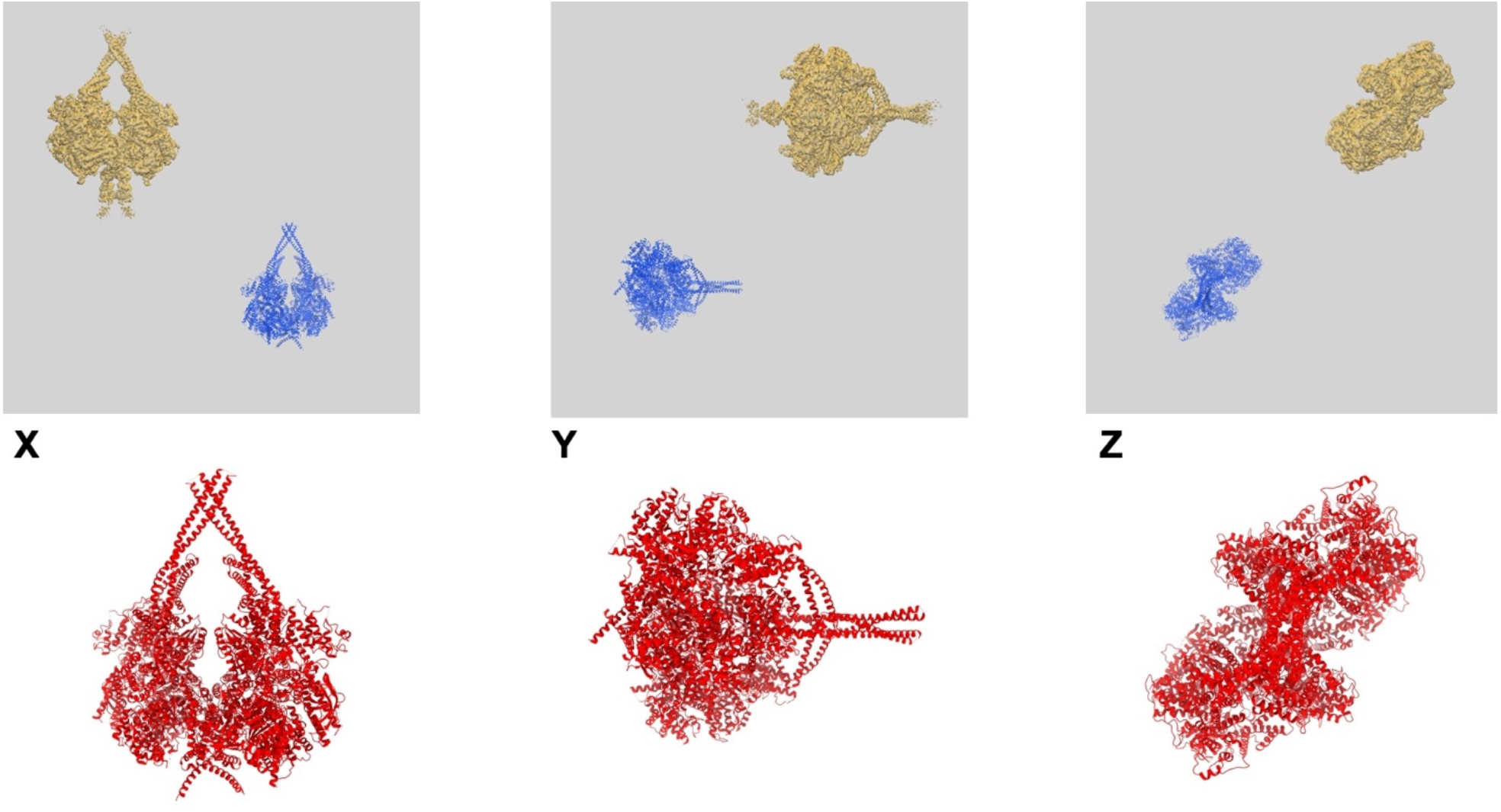
Example of a map-origin error, leading to a misaligned (and in this case, completely non-overlapping) map and model. The top row shows the map-model overlay views and the bottom row the model coloured by residue-inclusion score (viewed along the same axes), where the red colour of the entire model reflects the fact that not a single residue is inside the map.

**Figure 5** shows an example where there is a discrepancy in the relative scaling of map and model. Whereas the determination of the physical scaling of a model is an intrinsic part of the model-determination process, the same is not the case for SPA maps. It is not uncommon for errors in microscope-magnification calibration to propagate to the deposited voxel size leading to such scaling errors.

**Figure 5.**
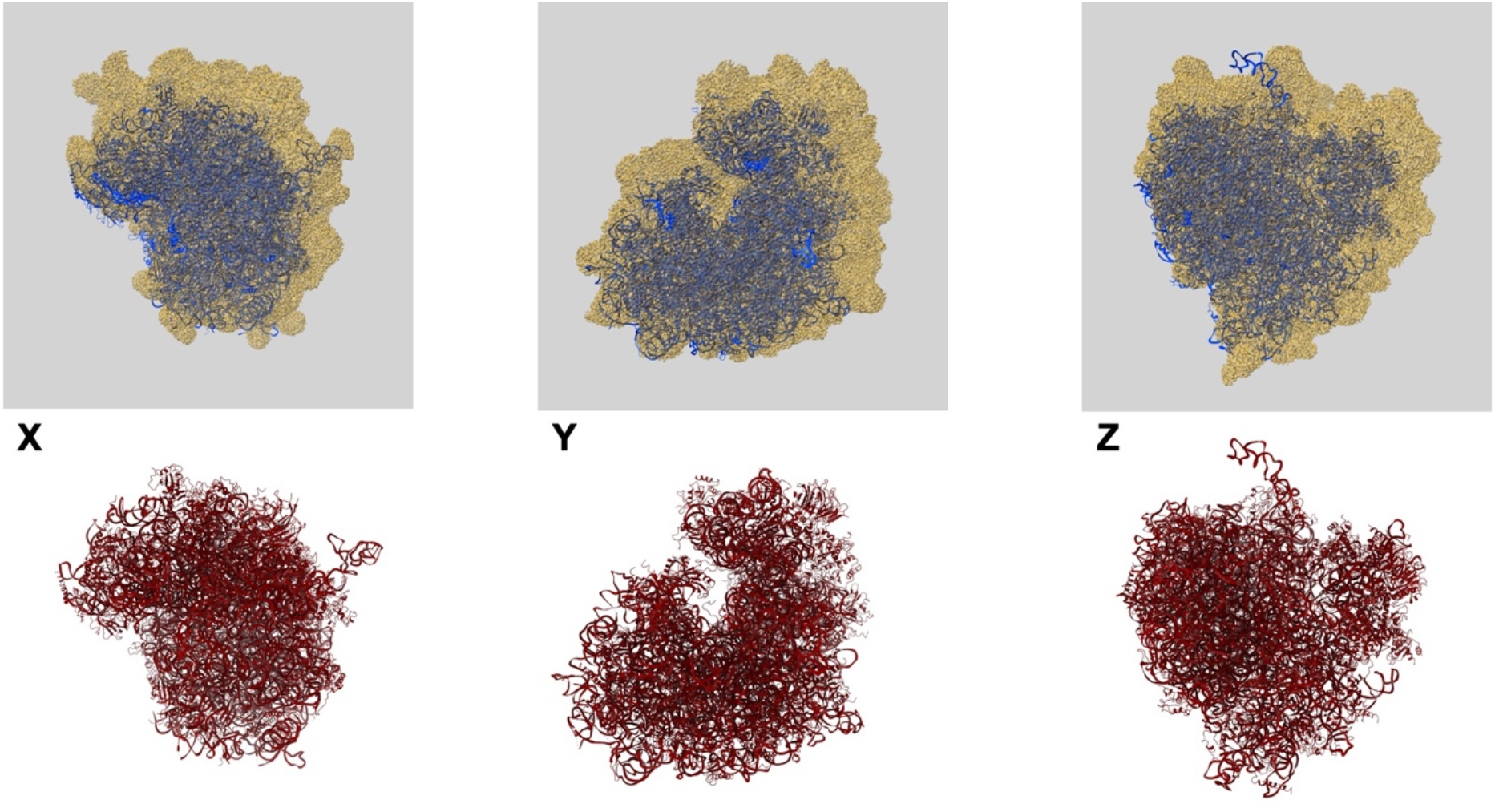
Example of a pixel-sampling error, leading to an incorrect relative scaling of map and model. The top row shows the map-model overlay views and the bottom row the model coloured by residue Q-score (viewed along the same axes).

Upon deposition to EMDB, the depositor provides the recommended contour level for viewing and rendering the primary map. It is also used to assess the fit of the model to the map using the atom and residue-inclusion scores. **Figure 6** shows what happens if the contour level selected is considerably too high or low. In panel (A) the contour level is set too high so that the map fails to cover the model properly, as reflected in the 3D view and the atom-inclusion plot (the map covers only ∼30% of the atoms). In panel (B) the contour level is set too low instead. This leads to a deceptively “good” atom-inclusion plot (100% of all atoms inside the map), but clearly the map looks unrealistic in the 3D view. The Q-score and 3D-Strudel measures are independent of the choice of contour level and hence do not suffer from inappropriate choices.

**Figure 6.**
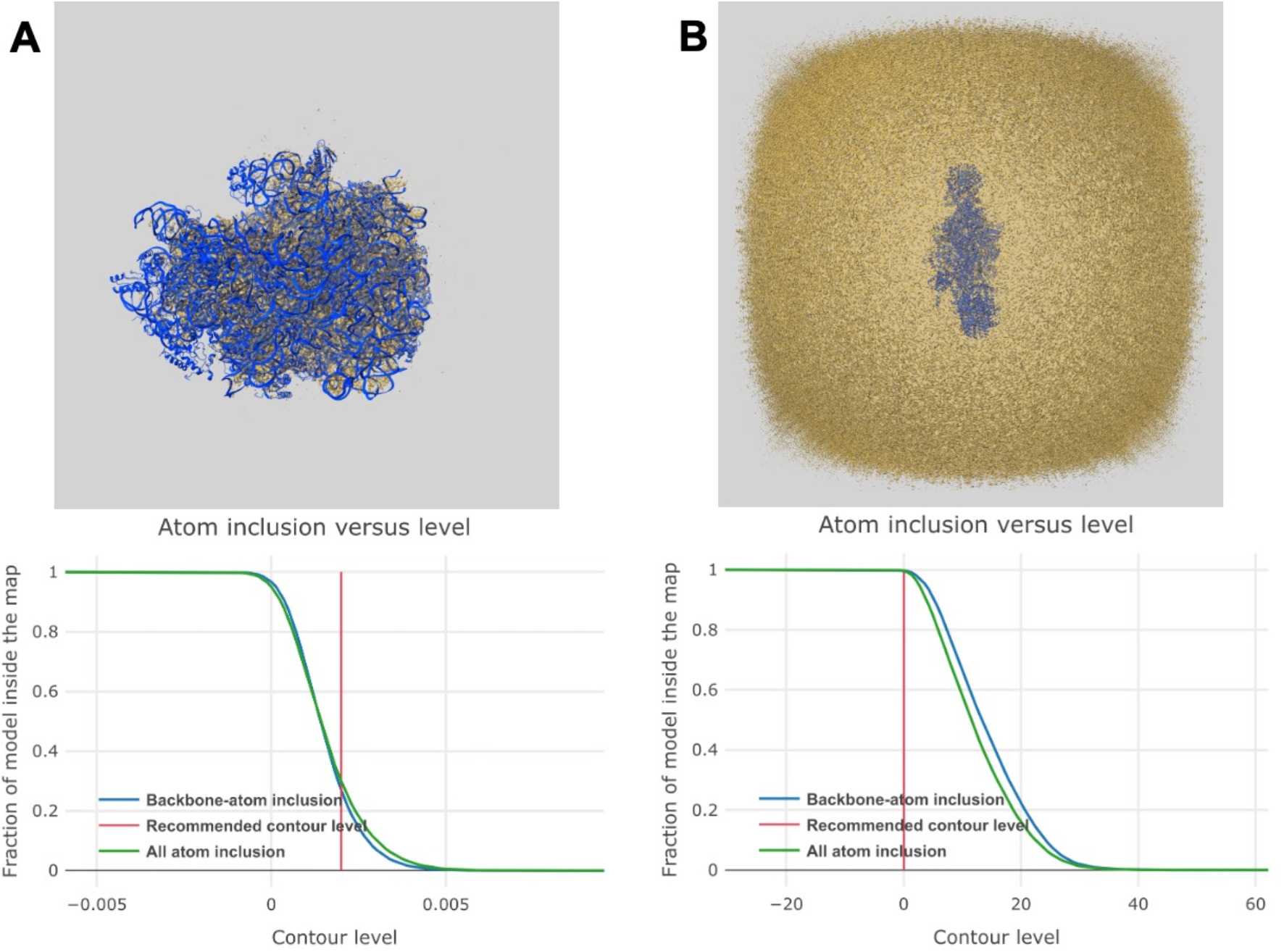
Effects of an inappropriate choice of contour level (see text for details). (A) The atom-inclusion graph and map-model overlay view for a case where the recommended contour level has been set too high. (B) The same components shown for a case where a contour level has been chosen that is too low.

**Figure 7** shows an example of two of the issues that can be identified with 3D-Strudel. In one case a phenylalanine residue has been modelled with an incorrect side-chain conformation that does not fit the map (also indicated by a low Q-score value of 0.31). In the other case, a stretch of residues has been built in a location where there is no support for them in the map. Again, low Q-scores (between - 0.18 and 0.27) confirm that there are issues with this part of the model. Both 3D-Strudel and Q-score are independent of contour level, so these are not issues that are caused by an incorrect choice of that level. 3D-Strudel can in addition be useful in detecting sequence-register errors between map and model (not shown; see Istrate *et al*., to be published).

**Figure 7.**
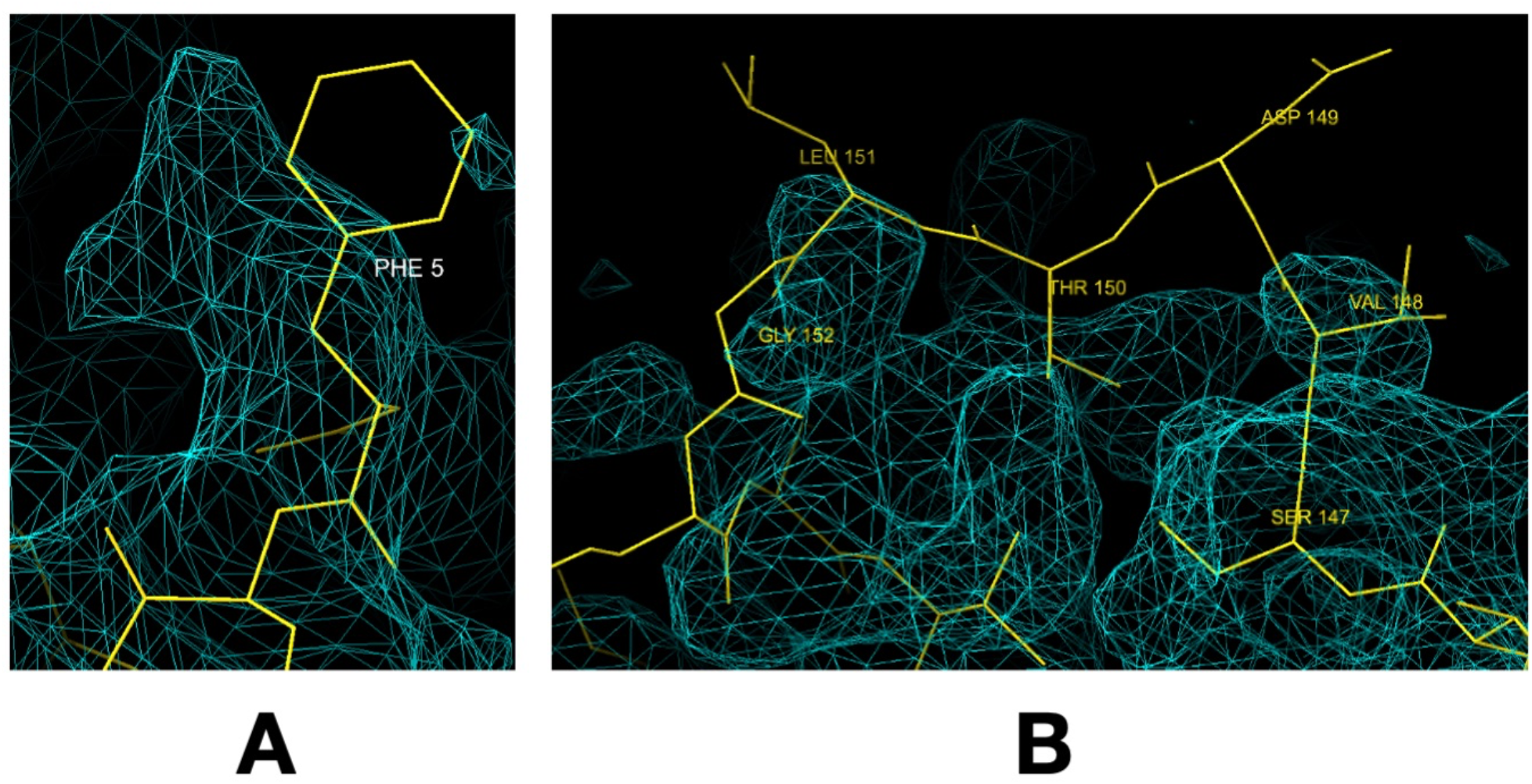
Examples of two map/model-fit issues highlighted by 3D-Strudel (see text for details). (A) A phenylalanine residue that has been modelled with an incorrect side-chain conformation. (B) A stretch of residues (147-153) that have been built in the absence of supporting map features.

## 4. Conclusions and plans

The primary goal of this work is to facilitate assessment and validation of cryo-EM data (maps, tomograms and models) by specialists and non-specialists alike, through the use of intuitive 2D and 3D visualisations and well-understood metrics, plots and graphs. This is achieved in particular with the validation tools in tiers 2 and 3 of our strategy. Tier 1 provides additional validation information suitable for specialists but also has a partly different purpose, namely to allow experimentation at scale with new validation approaches and metrics. For some validation tasks (e.g., assessment of local resolution or map/model-fit analysis) there are multiple competing methods whose relative merits are not yet understood. For example, we plan to integrate a number of tools to calculate local resolution, including ResMap (Kucukelbir *et al*., 2014), BlocRes (Bsoft) (Cardone *et al*., 2013), MonoRes (Vilas *et al*., 2018), and DeepRes (Ramírez-Aportela *et al*., 2019), as well as tools to assess anisotropy and angular coverage such as CryoEF (Naydenova & Russo, 2017) and MonoDIR (Vilas *et al*., 2020). Archive-wide analysis and comparison of such metrics, as well as inspection of results for individual structures by experts will hopefully bring more clarity and eventual community consensus about which tools are most appropriate to apply for a certain validation task. We are also developing a server with which the tier 1 pipeline can be run on data and models that are not yet in the public archives, and by incorporating the VA pipeline into the CCP-EM package (Wood *et al*., 2015) this can also be accomplished in-house. EMDB will continue to engage with developers of validation tools and include promising new methods into the VA pipeline.

The cryo-EM community has played a crucial role in this on-going project. We are grateful to all software providers who allow us to use their tools in the pipeline, and the input from the wwPDB single-particle cryo-EM data-management workshop (Kleywegt et al., to be published) has been invaluable in guiding our work on all three tiers of the validation resources. The workshop focussed exclusively on validation of molecular maps and models, and these are therefore covered most extensively at present. However, many of the 3D volume analyses are also applied to tomograms by all three tiers.

We expect that future improvements to EMDB and PDB metadata collection, more explicit file typing, mandatory deposition of certain types of data etc. will also improve our ability to validate cryo-EM structural data. Simultaneously, the field is still experiencing rapid growth and expansion as well as development of new or improved approaches to structure determination and analysis, many of which will impact the way in which we validate the data. We look forward to continuing to address all these challenges, in collaboration with the cryo-EM community.

## 5. Software dependencies and availability

The VA pipeline is a Python-based program that uses several standard Python packages (numpy, scipy). In addition, there are several external dependencies for EM-specific calculations, including CCP-EM (Wood *et al*., 2015; Burnley *et al*., 2017), TEMPy (Joseph *et al*., 2017), REFMAC (Murshudov *et al*., 1997), ProSHADE (Nicholls *et al*., 2018), EMDA (Warshamanage *et al*., 2021), 3D-Strudel (Istrate *et al*., to be published), Q-score (Pintilie *et al*., 2020) and ChimeraX (Goddard *et al*., 2018). Several other metrics have already been integrated but their results are not yet presented on the VA webpages. This includes CCC (Warshamanage *et al*., 2021), 3DFSC (Tan *et al*., 2017), SMOC (Joseph *et al*., 2016), and EMRinger (Barad *et al*., 2015), which in turn relies on cctbx (Grosse-Kunstleve *et al*., 2002).

The VA pipeline will be integrated into the CCP-EM package and the VA Python programming package is also available from PyPI (https://pypi.org/project/va/). Tier 3 validation is further accessible as part of the wwPDB validation resources through the OneDep validation server (https://www.wwpdb.org/validation/validation-servers) and the OneDep Python API (https://www.wwpdb.org/validation/onedep-validation-web-service-interface).

## Acknowledgments

We are grateful to our EMDB colleagues A. Istrate for many discussions, code reviews and support for 3D-Strudel integration, O. Salih and J. Turner for providing example applications, R. Pye and S. Abbott for integrating the tier 3 components into the wwPDB validation pipeline, and N. Fonseca for help with integrating the tier 2 components into the EMDB entry pages. We thank P. Korir and A. Iudin (EMPIAR) for technical discussions and J. Berrisford (PDBe) for help with the mmCIF format. We thank the wwPDB biocurators and partners for valuable feedback and bug reports.

We owe a debt of gratitude to all participants of the 2020 single-particle cryo-EM data-management workshop. Their feedback and recommendations have led to substantial improvements in the contents of all three tiers and indeed it was their suggestion to develop a three-tiered approach in the first place.

We thank the CCP-EM staff for software support and ongoing integration of the VA pipeline into CCP-EM. We further thank our collaborators in the Wellcome Trust-funded UK EM Validation Network for fruitful discussions: M. Topf and T. Cragnolini (also for help with TEMPy); G. Murshudov, R. Warshamanage and M. Tykac (also for help with REFMAC, EMDA, CCC and ProSHADE); P. Rosenthal, M. Winn, E. Orlova and A. Roseman. We thank Y.Z. Tan, P. Baldwin and D. Lyumkis for help with the implementation of the 3DFSC software, the PHENIX team for software support, J. Fraser and B. Barad for EMRinger support, G. Pintilie for help with the Q-score software, and T. Goddard and G. Couch for help with ChimeraX.

ZW is funded by the Wellcome Trust (grant 208398/Z/17/Z to P. Rosenthal). Work on EMDB at EMBL-EBI is further supported by the Wellcome Trust (grant 212977/Z/18/Z to AP and GK) and EMBL with funding from its member states.

